# Fast, accurate ranking of engineered proteins by receptor binding propensity using structure modeling

**DOI:** 10.1101/2023.01.11.523680

**Authors:** Xiaozhe Ding, Xinhong Chen, Erin E. Sullivan, Timothy F. Shay, Viviana Gradinaru

**Affiliations:** Division of Biology and Biological Engineering, California Institute of Technology, 1200 E California Blvd, Pasadena, 91125, CA, USA

**Keywords:** Protein engineering, AAV engineering, In silico screening, Receptor binding, Protein structure prediction

## Abstract

Deep learning-based methods for protein structure prediction have achieved unprecedented accuracy. However, the power of these tools to guide the engineering of protein-based therapeutics remains limited due to a gap between the ability to predict the structures of candidate proteins and the ability to assess which of those proteins are most likely to bind to a target receptor. Here we bridge this gap by introducing Automated Pairwise Peptide-Receptor AnalysIs for Screening Engineered proteins (APPRAISE), a method for predicting the receptor binding propensity of engineered proteins. After generating models of engineered proteins competing for binding to a target using an established structure-prediction tool such as AlphaFold-Multimer or ESM-Fold, APPRAISE performs a rapid (under 1 CPU second per model) scoring analysis that takes into account biophysical and geometrical constraints. As a proof-of-concept, we demonstrate that APPRAISE can accurately classify receptor-dependent vs. receptor-independent adeno-associated viral vectors and diverse classes of engineered proteins such as miniproteins targeting the SARS-CoV-2 spike, nanobodies targeting a G-protein-coupled receptor, and peptides that specifically bind to transferrin receptor or PD-L1. APPRAISE is accessible through a web-based notebook interface using Google Colaboratory (https://tiny.cc/APPRAISE). With its accuracy, interpretability, and generalizability, APPRAISE promises to expand the utility of protein structure prediction and accelerate protein engineering for biomedical applications.

## Introduction

Many protein-based biologics rely on precise targeting. As a result, protein engineers have devoted considerable effort to create specific binders, using methods such as directed evolution ^1–4^ and rational design ^5–7^. Currently, the costly experimental evaluation of candidate binders using *in vitro* and *in vivo* assays presents a bottleneck, which can be eased using computational prioritization ^8^.

Two strategies are employed to predict protein functions: end-to-end sequence-function and two-step sequence-structure/structure-function. End-to-end, sequence-function models can predict complex functions such as enzyme activities or ion channel conductivity ^9,10^, which are challenging to calculate using physical principles ^11^. However, such specialized models require domain-specific, high-quality training datasets for accurate prediction. In comparison, the two-step sequence-structure/structure-function strategy offers a more generalizable solution, particularly for functions with well-understood biophysical mechanisms such as protein-protein binding.

The rapid development of deep learning-based methods has brought unprecedented accuracy to the first step of the sequence-structure/structure-function strategy. Since AlphaFold2 (AF2)’s outstanding performance in CASP14 in 2020^12^, several new deep learning-based structure-prediction tools have been released ^13–24^, providing a diverse tool set for generating protein models with atomic-level precision. While the original AlphaFold2 can predict peptide-protein complexes ^25^, there are enhanced versions such as AlphaFold-Multimer that can model multi-chain complexes with greater accuracy ^14,17^. Importantly, these structure-prediction tools allow the generation of models in less than one GPU hour each, a level of throughput that experimental methods cannot match.

The second step, ranking target binding propensities based on structure predictions, has been less attended than the first. Structure-prediction tools generate confidence scores for predicted multimer models, such as pLDDT and pTM scores (used by AF2) ^12^, and interface pTM scores (used by AF-Multimer) ^14^, which have been used “off-label” as metrics to evaluate the probability of binding ^17,26^. However, previous reports ^27^ and our experience revealed that these scores alone are, in some cases, not reflective of binding propensities, particularly when the interaction is weak or transient. Extracting additional information stored in the 3D coordinates using biophysical principles may help improve the accuracy of binder ranking.

Ranking the binding probability of engineered proteins through modeled structures presents unique challenges. A frequent challenge is imposed by the high sequence similarity between candidate molecules. Engineered protein variants are often constructed by modifying a short variable region in a common scaffold. Due to this similarity, the energy difference between the candidate binders can be very small, sometimes buried in the error of the energy function used for candidate ranking ^28,29^. This problem is compounded by structure-prediction methods that rely heavily on co-evolutionary information or homology, causing them to generate similar binding poses for the candidate proteins. Another major challenge is assessing a large number of predicted structure models efficiently. Direct quantification of protein-protein interface energy using interpretable, physics-based methods trades off between accuracy and speed ^30^. For instance, molecular dynamics simulation methods can cost more than 10^3^ CPU hours per model. Faster, less rigorous methods with better-than-random ability to predict the impact of interface mutations still require 1 CPU minute to 1 CPU hour per non-antibody-antigen model ^30^. In the post-AlphaFold era, an interpretable and efficient method of predicting the target binding of a large number of models would greatly accelerate protein engineering efforts.

Recently, Chang and Perez utilized competitive modeling with AF-Multimer to demonstrate a correlation between competition results and peptide binding affinities ^27^. However, the study’s method of assessing the competition results necessitates a comparison of the modeled structures to an experimentally solved “native” structure, which is not available for many engineered proteins.

To bridge the remaining gap between structure prediction and protein engineering, here we present Automated Pairwise Peptide-Receptor AnalysIs for Screening Engineered proteins (APPRAISE), a readily interpretable and generalizable method for ranking the receptor binding propensity of engineered proteins based on competitive structure modeling and fast physics-informed structure analysis.

## Results

The workflow of APPRAISE (Figure 1) comprises four main components. In the first step, pairs of peptides from *N* candidate protein molecules (*N* ^2^ pairs total) are modeled in complex with a target receptor using a state-of-the-art structure method such as AF-Multimer ^14^. In the second stage, a simplified energetic binding score is calculated for each peptide (i.e., the peptide of interest (POI) and its competitor). In the third optional step, geometrical constraints for effective binding are applied to these scores. Finally, the result of each competition is decided using the score difference between the POI and the competitor, and the peptides are ranked based on the matrix of competition results.

**Fig. 1.**
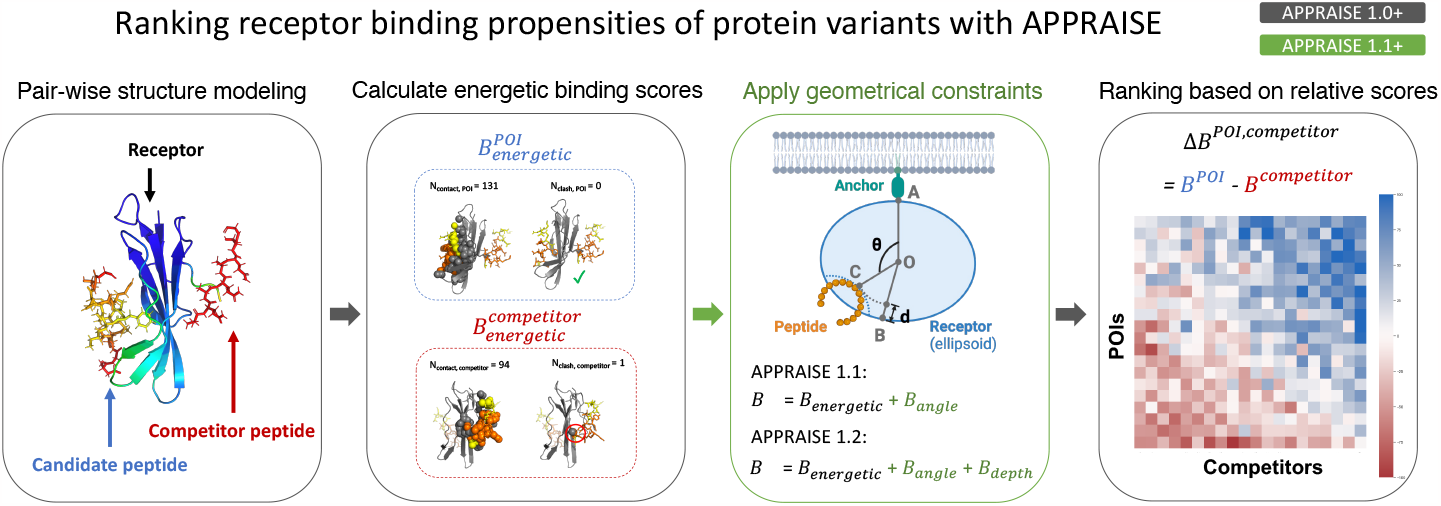
Workflow of Automated Pairwise Peptide-Receptor AnalysIs for Screening Engineered proteins (APPRAISE). First, peptides from the engineered protein candidates’ receptor-binding region are modeled in competing pairs with the target receptor using tools like AF-Multimer or ESMFold. Second, a non-negative energetic binding score based on atom counting is calculated for each peptide. Third, in APPRAISE 1.1+, additional geometrical constraints critical for peptide binding, including the binding angle and pocket depth, are considered. Finally, a relative score for each match is calculated by taking the difference between the scores for the two peptides. The averaged relative scores form a matrix that determines the final ranking.

### APPRAISE can accurately classify receptor-mediated brain transduction of viral vectors

We first developed APPRAISE to predict the binding propensities of engineered Adeno-Associated Viral (AAV) capsids for brain receptors. Recombinant AAVs are widely used as delivery vectors for gene therapy due to their relative safety as well as their broad and engineerable tropism. *In vivo* selections from libraries of randomized peptide-displaying AAV variants have yielded capsids that can transduce the animal brain^1,2,31–35^, an organ tightly protected by the blood-brain barrier (BBB). Widely-known examples among these capsids are AAV-PHP.B ^1^ and AAV-PHP.eB ^31^, two AAV9-based ^36^ variants displaying short (7-9 amino acids) surface peptides. The two variants can efficiently deliver genetic cargo to the brains of a subset of rodent strains. Genetic and biophysical studies have revealed that the BBB receptor for PHP.B/PHP.eB in these strains is LY6A, a GPI-anchored membrane protein ^37–39^. A dataset comprising peptide-displaying AAV capsids that were engineered in a similar way as PHP.B/eB was collected in order to train the APPRAISE method (Figure S1). Although binding between the AAV and the LY6A receptor is dynamic ^40,41^ and therefore challenging to quantitatively measure, we could infer the binary LY6A-binding profiles of AAV capsids from their differential brain transduction profiles in mouse strains with and without the receptor, producing a training set of peptide-displaying AAV capsids (Figure S1).

One challenge for modeling AAV capsids is that they are huge complexes made of 30,000+ amino acids (aa). In order to reduce computational costs for structure modeling and avoid complications arising from non-specific interactions, we modeled each AAV capsid variant using a single peptide spanning the engineered region (Figure 2a). This peptide (residues 587-594 in the VP1 sequence) includes 7 inserted residues and 8 contextual residues flanking the insertion. All of these residues are surface-exposed and may make direct contact with the receptor in the assembled capsid. Modeling this surface peptide (15 aa) is far less computationally intensive than modeling the entire capsid or even an asymmetric capsid subunit (500+ aa). In addition, compared to the latter, it may improve accuracy by eliminating competing interactions of residues normally buried in inter-subunit interfaces.

**Fig. 2.**
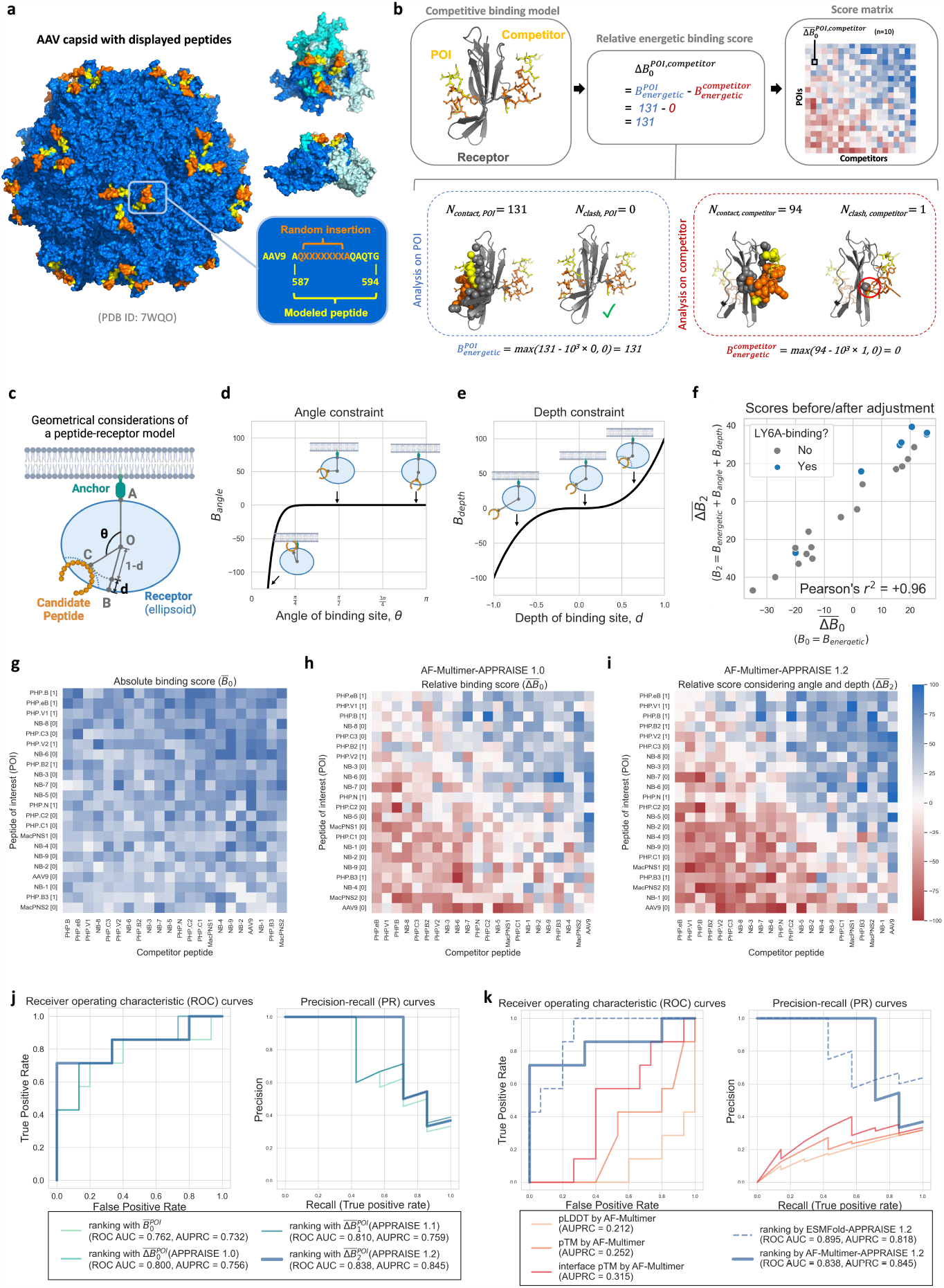
Binary classification of receptor-binding AAV capsids using physical and geometrical principles. **a**, a structure model of AAV-PHP.eB, highlighting the site for inserting the displayed peptide (orange) and the peptide used for APPRAISE modeling (yellow or orange). The left image shows the AAV capsid of 60 structurally identical subunits. The two images on the top right show a top view and a side view around the 3-fold axis, respectively. The three subunits that make the trimer are colored blue, cyan, and white. The sequence corresponding to the peptides is shown in the bottom right. **b**, An example showing the calculation process of a relative energetic binding score. The number of contacting atoms (*<* 5 Å) and the number of clashing atoms (*<* 1 Å) for each peptide in the competition are counted, and an absolute energetic binding score is calculated based on the counts according to Eq. 1. A difference between the two numbers, or the relative energetic binding score, is then calculated. The competition result between two peptides is determined using the average of relative binding scores across 10 replicates. The matrix of the mean scores is then used to rank the peptides of interest (POIs). **c**, A simplified geometrical representation of a peptide-receptor model, where the hull of the receptor is represented by an ellipsoid (blue). Point O: the center of mass of the receptor. Point A: the receptor’s terminus attached to an anchor. Segment OB: the minor axis of the ellipsoid receptor hull. Point C: the deepest point on the candidate peptide (orange). *θ*: the binding angle of the peptide. *d*: the binding pocket depth of the peptide. **d**, The angle constraint function. Three representative scenarios with different binding angles are highlighted. **e**, The depth constraint function. Three representative scenarios with different binding depths are highlighted. **f**, Comparison of the averaged relative binding energy scores before geometry-based adjustments vs. after adjustments. **g-i** Heatmaps representing the matrix of mean scores of 22 AAV9-based capsid variants, including **g** mean absolute binding scores, **h** mean relative binding scores, and **i** mean relative binding scores that have considered both angle and depth constraints. All heatmap matrices were sorted by point-based round-robin tournaments (Methods). Bracketed numbers in the row labels are LY6A-binding profiles of the capsids inferred from experimental evidence (Figure S1). Each block in the heatmap represents the mean score measured from 10 independent models. **j-k**, comparison of different ranking methods used as binary classifiers to predict the LY6A-binding profile of 22 AAV9-based capsid variants. **j**, comparison between rankings given by different versions of APPRAISE scores using AF-Multimer as the structure prediction tool. **k**, comparison between rankings given by confidence scores of AF-Multimer versus rankings given by APPRAISE 1.2 using either AF-Multimer or ESMFold as prediction engines. The sequence and shape parameters of LY6A used for the modeling and analyses are included in Table S1.

To discriminate relatively small differences in receptor binding propensities of candidate peptides, we modeled the peptides pairwise in competition for the target receptor^27,42^. To evaluate the competition results efficiently, we designed a score based on simple atom counting as a rough estimate of the interface free energy between the peptide of interest (POI) and the receptor in a structure model (Figure 2b). This score, which we term the energetic binding score (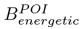, simplified as 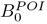), is a non-negative value calculated from the numbers of contacting and clashing atoms at the interface (Eq. 1). We describe the detailed rationale behind this score in the Methods.

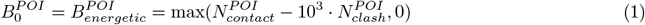

To take full advantage of the information encoded in the competitive models, we further derived a “relative binding score”, inspired by the “specificity strategy” for protein-protein interface design ^43^. The relative score takes the difference between the absolute scores for the POI and competitor peptide (Eq. 2), rewarding POIs destabilizing competing peptides’ binding.

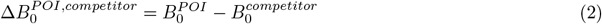

An engineered protein must meet certain geometrical constraints to effectively bind to a membrane receptor (Figure 2c). To utilize this geometrical information, which is likely unused by structure-prediction tools, we incorporated two essential constraints for effective binding: the binding angle and the binding depth (Figure 2c-e).

The first constraint comes from the angle a binding protein can make (Figure 2c,d). In modeling a peptide-receptor complex using the extracellular domain of the membrane receptor (e.g., LY6A), most structure predictors (e.g., AF-Multimer) would consider the whole surface of the domain to be accessible by the peptide. However, in biological conditions, the membrane-facing side of the receptor is inaccessible to the engineered peptide. This polarity of accessibility is a general property of any receptor that is closely anchored to a larger complex. To account for the potentially huge energy cost of an engineered peptide binding these inaccessible locations, we used a steep polynomial term to penalize peptides that bind to the anchor-facing part of the receptor (Figure 2d, defined in the Methods by Eq. 5). 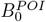 is adjusted by this geometrical constraint term, rectified to be non-negative, and 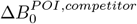 is also re-calculated accordingly, yielding new scores 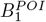 and 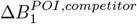 (Eq. 3).

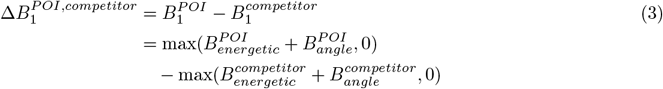

The second constraint concerns the binding pocket depth (Figure 2c,e). We hypothesized that peptides binding to a deeper pocket on the receptor surface might benefit from longer receptor residence time, which is vital for the efficacy of many therapeutics ^44^. Based on this hypothesis, we included a pocket depth consideration in APPRAISE’s scoring function. We used a relative pocket depth measurement instead of an absolute peptide-receptor distance measurement to avoid possible bias caused by the sizes of different receptors. We then used an odd polynomial term to reward peptides that insert into deep pockets on the receptor while penalizing peptides that attach to surface humps (Figure 2e, defined in the Methods by Eq. 6). The addition of the depth term gives us an adjusted score 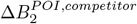 (Eq. 4).

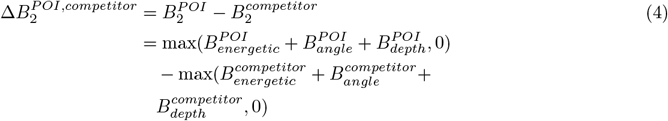

We compared different versions of scoring methods based on competitive modeling results using AF-Multimer modeling (Figure 2f-i). Individual matching scores with statistical significance were used to determine wins and losses, and the total matching points in a tournament were used to rank all candidate proteins (Methods). We found that simple atom-counting-based 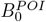 can already differentiate LY6A-binding peptides from non-binders (Figure 2g, j). Compared to 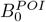 alone, the relative score 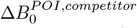 showed improved prediction power, a ROC AUC of 0.800 and an AUPRC of 0.756 for the training dataset (Figure 2h, j, k). Adding both geometrical terms, *B*_*angle*_ and *B*_*depth*_, into consideration indeed improved the prediction accuracy of the binding score (Figure 2f, i-k), yielding a ROC AUC of 0.838 and an AUPRC of 0.845 (Figure 2j, k). Importantly, the improvement in ROC AUC mainly came from the low-false-positive-rate segment of the ROC curve, which is crucial for *in silico* screening of engineered proteins. For clarity, we name the version that considers only the angle constraint (through score Δ*B*_1_) APPRAISE 1.1 (Figure S2a) and the version that considers both angle and depth constraints (through score Δ*B*_2_) APPRAISE 1.2 (Figure 2i).

We then compared AF-Multimer-based APPRAISE 1.2 with other structure-based peptide affinity ranking methods on the AAV dataset (Figure 2k). With this particular dataset, the model confidence scores pLDDT, pTM, and interface pTM failed to differentiate whether an AAV variant is an LY6A binder, producing worse-than-random prediction (ROC AUC *<* 0.5). This is possibly due to the dynamic nature of the interaction between LY6A-binding AAV variants and the receptor ^40,41^, which causes the confidence scores of the complex models to be generally low. APPRAISE 1.2 utilizing ESMFold as the structure prediction engine, however, performed at a comparable level to AF-Multimer-APPRAISE 1.2 (Figure S2b), with a ROC AUC of 0.895 and AUPRC of 0.818 (Figure 2k).

AF-Multimer-APPRAISE 1.2 ranking outperformed all other ranking methods at the low-false-positive-rate end of the ROC curve, with a true positive rate of 0.714 and no false positive predictions. The performance with stringent cut-off values is particularly relevant for protein engineering applications, where the goal is typically to identify a few positive binders from many negative, non-binding candidates. The superiority of AF-Multimer-APPRAISE 1.2 in dealing with this kind of imbalanced library is also evidenced by its highest AUPRC. Because of this, we chose to characterize AF-Multimer-APPRAISE 1.2 further. In the following text, ‘APPRAISE’ will be used to refer to AF-Multimer-APPRAISE 1.2 unless otherwise specified.

### APPRAISE is generally applicable to diverse classes of engineered proteins

To determine the applicability of APPRAISE to different classes of engineered proteins, we applied the method to four classes of engineered protein binders targeting four representative receptors for therapeutics.

We first applied APPRAISE 1.2 to other short peptide binders (Figure 3a-d). In the first trial, the method successfully ranked a peptide selected by phage display to bind human transferrin receptor^4^, a well-characterized BBB receptor, over non-binding counterparts from the same selection ^4^ (Figure 3a). In the second trial we evaluated two 47-aa-long, rationally-designed PD-L1 binding peptides ^7^ against the scaffold and length-matched AAV variable region fragments. Both designed PD-L1 binding peptides were clear winners, with the higher-affinity MOPD-1 peptide topping the list despite a high degree of sequence similarity (Figure 3c, Figure S3a).

**Fig. 3.**
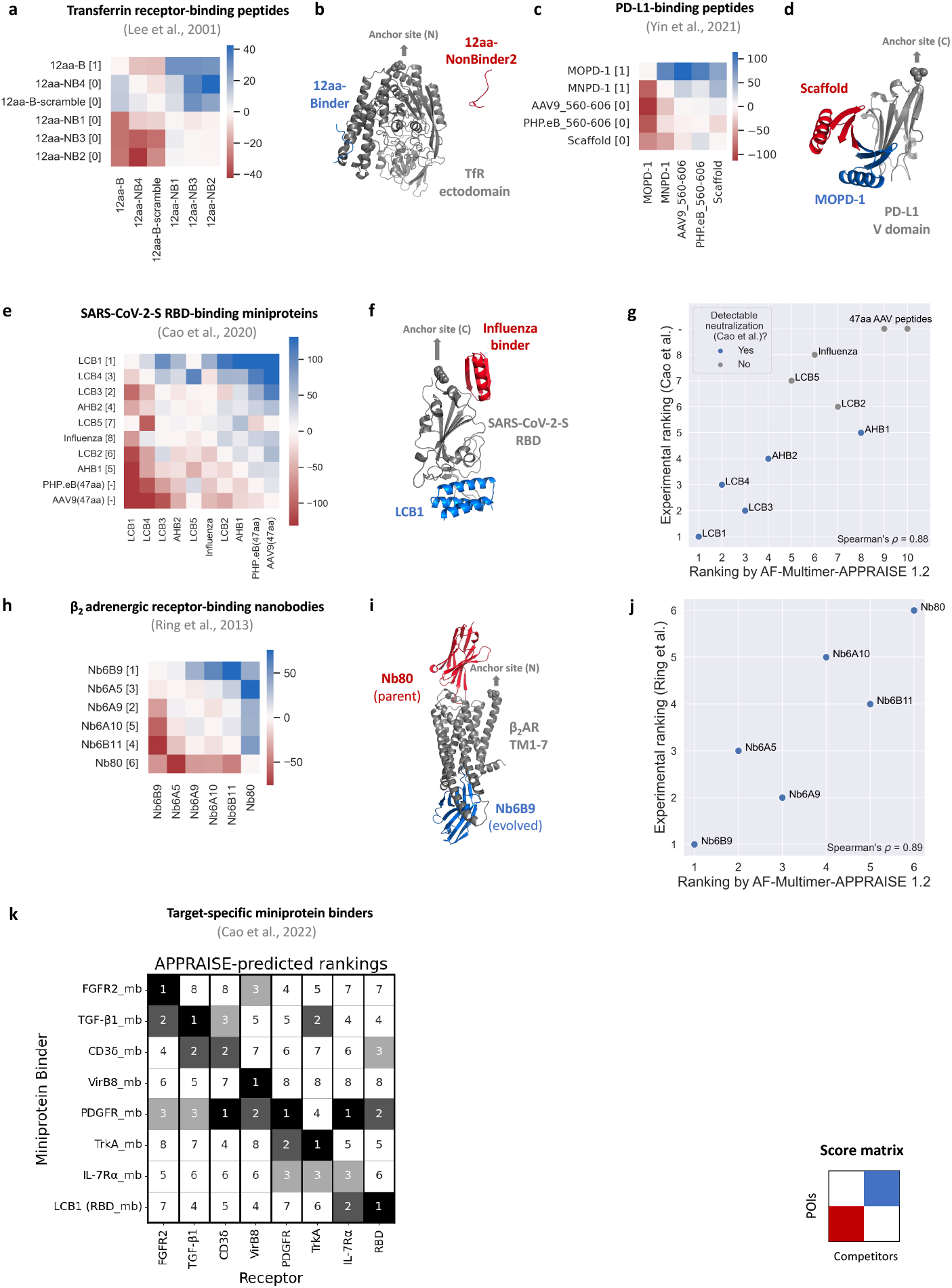
AF-Multimer-APPRAISE 1.2 accurately ranks binding propensities of different classes of engineered proteins. **a-b**, APPRAISE 1.2 ranking of transferrin receptor-binding peptides and non-binding control peptides ^4^. **a**, Pairwise score matrix and ranking of a panel of 12-aa peptides given by APPRAISE 1.2. Bracketed numbers in the row labels are experimentally determined transferrin receptor-binding profiles of each peptide ^4^. **b**, A representative AF-Multimer model result of a binding peptide (blue) competing against a non-binding peptide (red) for binding to transferrin receptor. **c-d**, APPRAISE 1.2 ranking of PD-L1-binding peptides and non-binding control peptides ^7^. **c**, Pairwise score matrix and ranking of a panel of 47-aa peptides given by APPRAISE 1.2. Bracketed numbers in the row labels show the PD-L1-binding profile of each peptide determined either experimentally (for MOPD-1, MNPD-1, and scaffold protein) or by expectation (for AAV9 and PHP.eB) ^7^. **d**, A representative AF-Multimer model result of MOPD-1 (blue), a designed binding peptide, competing against a non-binding scaffold peptide (red) for binding to PD-L1. **e-g**, APPRAISE 1.2 ranking of SARS-CoV-2-S RBD-binding miniproteins ^5^. **e**, Pairwise score matrix and ranking given by APPRAISE 1.2. Bracketed rankings in the row labels are determined based on experimentally-measured IC50 of each miniprotein to neutralize live SARS-CoV-2 ^5^. **f**, A representative AF-Multimer model result of LCB1 (blue), a SARS-CoV-2-S RBD-binding miniprotein, competing against an influenza virus-binding miniprotein ^6^ (red). **g**, A scatter plot showing the correlation between APPRAISE-predicted ranking and experimentally-measured IC50 ranking of all miniproteins tested. Blue points highlight binders that showed the capability of complete neutralization of the SARS-CoV-2 virus in the tested range of concentration *in vitro*. **h-j**, APPRAISE 1.2 ranking of *β*_2_ adrenergic receptor-binding nanobodies ^3^. **h**, Pairwise score matrix and ranking given by APPRAISE 1.2. Bracketed numbers in the row labels are rankings of experimentally-measured binding of each nanobody ^3^. **i**, A representative AF-Multimer model result of Nb6B9 (blue), the strongest evolved binder to active *β*_2_AR, competing against Nb80 (red), the nanobody used for the evolution. **j**, A scatter plot showing the correlation between APPRAISE-predicted ranking and experimentally-measured ranking by *β*_2_AR binding of all nanobodies tested. Each block in the heatmap represents the mean score measured from 10 independent models. For comparison, rankings given by AF-Multimer-APPRAISE 1.0, ESMFold-APPRAISE 1.2, and interface pTM of SARS-Cov2-S RBD-binding miniproteins and *β*_2_ adrenergic receptor-binding nanobodies are shown in Figure S4. **k**, A summary of APPRAISE rankings of eight miniproteins ^45^ designed to bind to eight different target receptors. Figure S5 displays the score matrices utilized for rankings with individual receptors. Tables S1 and S2 include sequences and shape parameters of all receptors.

We next tested whether APPRAISE 1.2 can be used to evaluate larger proteins, for example, computationally designed miniproteins (50-90 aa) that bind to the receptor-binding domain (RBD) of SARS-CoV-2 spike protein^5^ (Figure 3e-g). Among the designed miniproteins, 5 can neutralize live SARS-CoV-2 virus *in vitro* with IC50 from 20 pM to 40 nM ^5^. The APPRAISE 1.2 rankings of the 5 neutralizing miniproteins matched well with their IC50 rankings (Spearman’s *ρ* = 0.90, *p* = 0.037, Figure 3g). The predictive accuracy of APPRAISE decreased when non-neutralizing miniproteins ^5^ and control AAV fragments were included (Spearman’s *ρ* = 0.88, *p <* 0.001, Figure 3g); nevertheless, the top 4 binders still remained on the top. In contrast, the ranking given by the iPTM score of AF-Multimer only achieved a Spearman’s *ρ* of 0.67 (*p* = 0.035).

We also used APPRAISE to rank 6 nanobodies (120 aa) that were evolved experimentally ^3^ with highly similar scaffolds (Figure S3b) to bind to an activated conformation of *β*_2_ adrenergic receptor (*β*_2_AR), a G-protein-coupled receptor (GPCR) (Figure 3h-j). APPRAISE 1.2 correctly found the strongest evolved binder and placed the parent (the weakest binder among all candidates) at the bottom (Figure 3h). The overall predicted ranking correlated well with the ranking from experimentally determined binding readouts^3^ (Spearman’s *ρ* = 0.89, *p* = 0.02, Figure 3j), surpassing the prediction given by the iPTM score of AF-Multimer (Spearman’s *ρ* = 0.49, *p* = 0.329, Figure S4f). The ability of APPRAISE to rank the binding affinity of nanobodies, a widely-used therapeutic modality, has the potential to expand its value in drug design and development. However, predicting the structure of larger adaptive immune complexes like IgG-antigen complexes is still considered a difficult task in general, and further improvements in the underlying structure prediction methods that APPRAISE relies on are required to generalize the ranking capability to these targets.

To evaluate the cross-receptor capabilities of APPRAISE, we used the method to rank eight recently developed miniproteins binding eight different therapeutically significant receptors ^45^. This ranking included all receptors with a ligand binding domain that is smaller than 250 aa in the Cao et al. study. APPRAISE accurately identified the correct binder within the top 3 in every instance, and 6 out of 8 times, the correct binder was ranked as the top 1 (Figure 3k, Figure S5).

We next compared the performance of AF-Multimer-APPRAISE 1.2 to alternative methods on both the miniprotein dataset and the nanobody datasets. AF-Multimer-APPRAISE 1.2 again yielded the most accurate predictions when compared to AF-Multimer-APPRAISE 1.0, ESMFold-APPRAISE 1.2, or interface pTM scores given by AF-Multimer (Figure S4), reflected by higher Spearman’s correlation to experimental rankings. ESMFold-APPRAISE 1.2 failed completely with the miniprotein dataset (Figure S4b). Upon further inspection, we found that the unfolded SARS-Cov-2-S RBD structure in ESMFold-generated complex models can explain the failed ranking prediction.

Without any fine-tuning, AF-Multimer-APPRAISE 1.2 demonstrated consistent prediction ability for ranking all four classes of proteins, including experimentally-selected and rationally-designed peptides, computationally-designed miniproteins, and nanobodies. Realizing the potential general applicability of the APPRAISE method, we have created a web-based notebook interface to make it readily accessible to the protein engineering community (Figure S6).

### HT-APPRAISE screening can identify novel receptor-dependent capsid variants

We next adapted APPRAISE 1.2 for *in silico* screening. The computational cost in the pairwise competition mode grows quadratically with the number of input variants, which is unsuitable for high-throughput screening. To address this scalability issue, we designed a two-stage screening strategy named high-throughput (HT)-APPRAISE (Figure 4a). The first stage aims to shrink the size of the variant library using a less accurate yet more scalable strategy. Variants are randomly pooled into groups and compete for receptor binding. The variants are then ranked by their absolute score 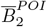. The number of pooled competitions grows linearly with the number of variants in the starting library, making the first stage of HT-APPRAISE suitable for larger libraries. In the second stage, the top variants selected from the first stage compete pairwise, yielding a matrix of 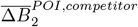 and a more accurate ranking.

**Fig. 4.**
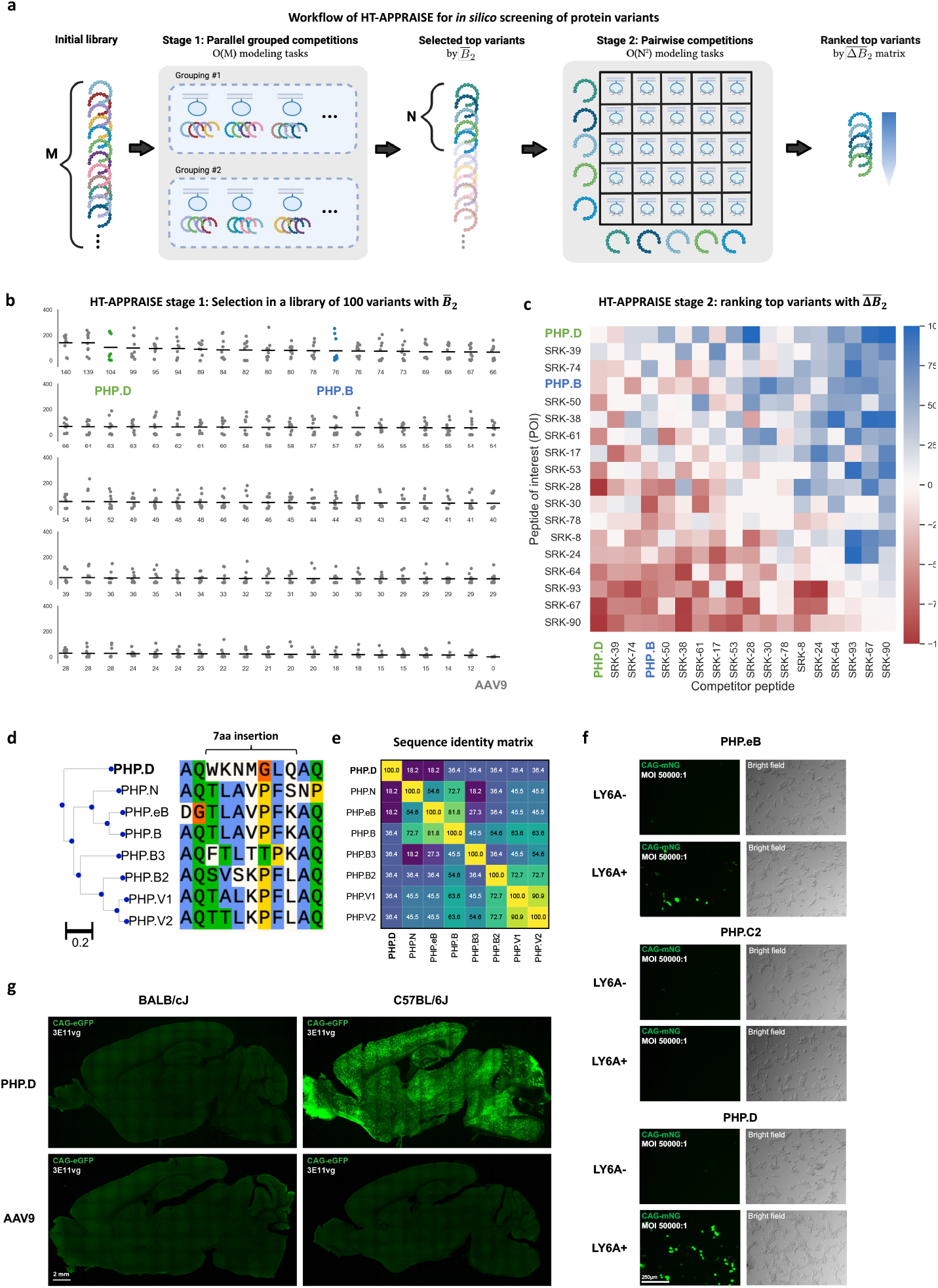
An *in silico* HT-APPRAISE screening of a medium-sized AAV library identifies a LY6A-dependent variant with a distinct sequence. **a**, A schematic showing the two-stage strategy for *in silico* screening of a variant library. In the first stage, M variants of interest are randomly pooled into groups of 4 and compete for receptor binding. At least two parallel groupings are used to reduce bias. Each peptide’s mean absolute binding score in the pool competitions is used for selecting the top N variants. In the second stage, the top N variants compete pairwise using standard APPRAISE for a more accurate ranking. **b-c**, Results from a proof-principle screening with 100 AAV9-based variants, including the wild-type control and variants with 7 aa insertions. Using a standard random algorithm, a total of 97 variants were picked from a list of 9000 variants ^2^ that demonstrated higher brain enrichment than the wild-type AAV9 after one round of screening in C57BL/6J mice (“Round 2 library” in ^2^). PHP.B and PHP.D, two known brain-transducing capsids, and wild-type AAV9, are spiked into the library. Table S4 shows the peptide sequences used in the screening. **b**, Stage 1 result. Dots indicate absolute binding scores measured from individual structure models. Horizontal bars indicate the mean scores of each variant. Scores of PHP.D (ranked 3^*rd*^), PHP.B (ranked 13^*th*^), and AAV9 (ranked 100^*th*^) are highlighted. **c**, Stage 2 result. Rows corresponding to scores of PHP.D (ranked 1^*st*^) and PHP.B (ranked 4^*th*^) are highlighted. Each block in the heatmap represents the mean score measured from 10 independent models. **d-g**, Characterization of PHP.D, a variant that tops the *in silico* screening. **d**, Sequence alignment and phylogenetic tree of known LY6A-dependent brain transducing variants. The sequence of PHP.D is very different from all other variants. The alignment and sequence distances were generated with Clustal Omega ^46^. The colored alignment is plotted with Snapgene software. Blue: conserved hydrophobic residues; green: conserved hydrophilic residues; orange or yellow: conserved unique residues (glycine or proline). **e**, Sequence identity matrix of the LY6A-dependent variants. **f**, *In vitro* infectivity assay in HEK293T cells. PHP.D and PHP.eB showed LY6A-enhanced transduction, while the negative control PHP.C2 did not show LY6A-enhanced transduction. AAV capsids carrying a fluorescent protein expression cassette were applied to HEK293T cells either transfected with LY6A or not at 5 × 10^8^ vg per well in a 96-well plate. Images were taken 24hr after transduction. n=3 per condition. Scale bar, 250 μm **g**, *In vivo* brain transduction of PHP.D vs. AAV9 in two mice strains. PHP.D showed transduction only in the LY6A+ strain, C57BL/6J. AAVs carrying CAG-mNeonGreen transgene were injected intravenously at 3 × 10^11^ vg per animal, and the tissues were harvested and imaged 3 weeks after injection. n=3 per condition. Scale bar, 2 mm.

We used our HT-APPRAISE *in silico* screening to find LY6A binders in a library of 100 capsid variants (Figure 4b-c). This library is composed of 97 capsid variants randomly chosen from a list of 9,000 variants showing superior brain enrichment in C57BL/6J mice compared to the wild-type AAV9 capsid^2^ as well as three spiked-in capsids: the variants PHP.B and PHP.D, and the wild-type AAV9 (Table S4). PHP.D is a brain-transducing capsid identified in a recent directed evolution campaign in our lab, the relevant receptor for which was unknown.

The HT-APPRAISE screening took 24 hours using 3 parallel Google Colaboratory GPU sessions and a laptop computer. In both stages of the HT-APPRAISE, the most time-consuming step was the structural prediction, which took approximately 0.1-1 GPU minute per peptide-LY6A model (a complex made of 114 aa total). In comparison, the time cost for structural analysis was negligible, taking less than 1 second per model on a CPU.

After the first stage of screening, both PHP.D and PHP.B appeared in the top 15% of the library (ranked by 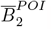) (Figure 4b).). In the second stage, the top 18 capsids were ranked using pairwise APPRAISE 1.2 (Figure 4c). PHP.D and PHP.B were the 1^*st*^ and the 4^*th*^ in the final ranking.

The most intriguing aspect of PHP.D’s result is that its variable region bears little sequence similarity to any of the LY6A-dependent variants used to develop the APPRAISE method (Figure 4d-e). To confirm this prediction result, we experimentally tested PHP.D’s LY6A dependency. An *in vitro* viral infection assay showed that PHP.D indeed exhibits LY6A-enhanced transduction of HEK293T cells (Figure 4f). In addition, *in vivo* systemic delivery of PHP.D packaging a ubiquitously-expressed fluorescent protein revealed that the brain transduction capability of this capsid variant is restricted to LY6A-expressing mouse strains (Figure 4g). The ability of HT-APPRAISE to identify binders with distinct sequences highlights the generalizability of the physics-informed, sequence-structure/structure-function strategy.

## Discussion

Here we describe APPRAISE, a structure-based, physics-informed method that accurately ranks receptor binding propensities of engineered proteins. APPRAISE uses a competitive, pairwise modeling strategy to capture affinity differences between even proteins with similar sequences and takes into account biophysical and geometrical principles.

The competition-based structure modeling strategy addresses the challenge of assessing small differences in binding affinity with high accuracy. This challenge was highlighted by a recent benchmarking study using AF2 models for molecular docking of small-molecule antibiotic candidates. The authors reported that the prediction power of the direct physics-based scoring is no better than a random model^47^. By contrast, a competition-based modeling strategy might have helped cancel the shared noise and amplify the signal arising from the small affinity differences. The competition setup, in many cases, forces the structure-prediction neural network to put only the more probable binder close to the receptor (Figure 3 d, i), converting a small probabilistic difference into a binary output.

The generalizability of AF-Multimer-APPRAISE is shown by its accurate ranking of five different classes of engineered proteins and twelve different receptors. This generalizability may be grounded in the physical principles learned by AlphaFold ^27,48^. For example, consistent with the report of ^27^, we also observed a recurring trend in our tests with AF-Multimer where binders ranked at the top are frequently predicted with secondary structures, which may indicate a stable binding interface. While we were revising this manuscript, APPRAISE has been applied in various scenarios, extending beyond the examples presented in this manuscript. For instance, in a recent study uncovering previously unknown blood-brain barrier receptors used by engineered AAVs, APPRAISE helped classify the AAVs based on their receptor specificity ^49^.

A key feature of APPRAISE is that its analysis module uses only information stored in the 3D coordinates, making the modular pipeline compatible with other computational tools for protein engineering. For example, the structure-prediction tool used in APPRAISE can be replaced by any current or future structure-prediction tool. Moreover, APPRAISE can be an orthogonal validation tool for structure-based protein design methods^50–52^, particularly those that rely on optimization of predicted confidence scores^53^.

The scalable, two-stage HT-APPRAISE strategy we designed allows *in silico* screening of protein candidates for receptor binding (Figure 4). Such screening can help prioritize leading candidates during drug discovery, reducing the huge time, financial, and environmental costs of experimental validation. For example, the computational tasks needed to screen 100 AAV variants that we presented here (Figure 4) could be completed within 24hr with research-grade computational resources. *In vivo* characterization of the capsids at a comparable scale would have taken several months.

As a competition-based ranking method, APPRAISE faces several intrinsic limitations. One such limitation is that APPRAISE only outputs the relative, not the absolute, probability of binding. Therefore, unless there are positive controls with known binding to compare against, a variant’s position at the top of the ranking does not indicate that the variant has an experimentally detectable binding affinity. Another limitation lies in APPRAISE’s assumption that the binding of competing proteins is mutually exclusive. Counterexamples arise if the competing proteins exhibit cooperative binding or attach to epitopes situated at a considerable distance. Furthermore, certain candidate proteins may exhibit a tendency to interact with one another rather than with the designated receptor. Additionally, the geometrical scores utilized in APPRAISE 1.1+ were computed assuming the receptor has a predominantly convex structure. Thus, these scores are most effective when applied to single protein domains with convex shapes.

Other limitations of APPRAISE may arise from the protein structure prediction engine that it relies on. For example, ESMFold-APPRAISE fails when the language-model-based structure prediction tool cannot properly fold the protein in a complex (Figure S4b). At the same time, AF-Multimer-APPRAISE results can be biased by the specific selection of multiple-sequence alignments due to the dependence on co-evolutionary information by AF-Multimer. So far, the APPRAISE pipeline’s ability to accurately rank IgG antibodies is limited due to the complexity of predicting antibody-antigen complex structures. Moreover, the accuracy and speed of APPRAISE may be compromised when the modeled proteins contain long disordered regions or large domains that are unnecessary for binding. As a result, pre-screening of several truncated protein constructs for minimal folding domains with the particular structure prediction tool (analogous to the common practice in structural biology) is always helpful. Additionally, the APPRAISE method is ineffective in ranking weak binders in a pool (e.g., Figure 3g), perhaps because the predicted structures do not offer many opportunities for meaningful interaction, resulting in near-zero competition scores. Fortunately, this should not be a practical concern for most protein engineering applications since the most valuable candidates usually bind with higher affinities. Considering these limitations, it is essential to conduct spot checks on model results to confirm their physical soundness.

While APPRAISE has succeeded in ranking the binding propensities of different protein variants, its accuracy and speed could be further improved. For example, the parameters of APPRAISE 1.2’s scoring function have only been minimally tuned to proper orders of magnitude to decrease the risk of over-fitting (Methods). With further fine-tuning of parameters and the ever-growing power of protein structure prediction, the APPRAISE method promises to streamline the process of engineering protein-based therapeutics.

## Methods

### Structure Modeling

#### Modeling of peptide-receptor complexes using AF-Multimer

Peptide-receptor models are modeled using Colabfold (Python package index: alphafold-colabfold 2.1.14), an implementation of integrated multiple-sequence alignment generation with MMseqs2 and structure modeling with AF-Multimer-v2^14,15,54–57^. First, batches of *.fasta files containing combined receptor sequences (Tables S1 and S2) and peptide sequences for the pairwise competition or pooled competition, where the “:” symbol separates the protein chains, are prepared using the function (appraise.input fasta prep.get complex fasta()). Second, the *.fasta files are used as input files for the “batch” Jupyter notebook in the Colabfold package, and the notebook is run on Google Colaboratory using a V100 SXM2 16GB GPU or an A100 SXM4 40GB GPU. The settings used for the modeling are listed below:

`msa_mode = “MMseqs2 (UniRef+Environmental)”`

`num_models = 5`

`num_recycles = 3`

`stop_at_score = 100`

`use_custom_msa = False`

`use_amber = False`

`use_templates = True`

`model_type = “auto” #or “alphafold2_multimer_v2”`

#### Modeling of peptide-receptor complexes using ESMFold

To model the peptide-receptor complexes using ESMFold, a process analogous to the one employed for AF-Multimer modeling is implemented. First, batches of *.fasta files containing combined receptor sequences (Tables S1 and S2) and peptide sequences for the pairwise competition or pooled competition, where the protein chains are separated by a poly-glycine linker (30 glycine residues), are prepared using the same Python function(get complex fasta()) mentioned above. Second, the *.fasta files are used as input files for a custom Jupyter notebook with codes adapted from the Colabfold package for batch modeling using ESMFold, and the notebook is run on Google Colaboratory using an A100 SXM4 40GB GPU. The custom Colab notebook is included in the APPRAISE package:

`appraise/misc_utilities/ColabFold_ESMFold_batch_run.ipynb`.

### Physics-informed analysis of individual structure models

The output folder containing *.pdb files generated by alphafold-colabfold is downloaded to a local computer for processing. Key parameters in a predicted structure model are measured, and the measurements are used to generate binding scores for each peptide in a model.

#### Automated quantification of the peptide-receptor models

The structure models are analyzed using PyMOL 2.3.3 using a custom PyMOL script. Briefly, the script loads all *.pdb models in a directory, extracts metadata from the file names, and measures the relevant contact atom numbers, angles, and distances. The measurements are saved as a *.csv file. The custom PyMOL script is included in the APPRAISE package:

`appraise/pymol_quantify_peptide_binding.py`.

#### Measurement of the *R*_*minor*_ of the receptor hull

The receptor shape parameter *R*_*minor*_, which is necessary for APPRAISE 1.2, is obtained by measuring the shape parameters of an AlphaFold-modeled receptor structure. Briefly, the monomeric receptor (Table S1) is modeled using Colabfold (Python package index: alphafold-colabfold 2.1.14). The top model is then analyzed using HullRad v8.1^58^ to obtain its major axis diameter *D*_*max*_ and aspect ratio *P*. *R*_*minor*_ is then calculated using the formula *R*_*minor*_ = *D*_*max*_*/P/*2. Before the analysis, *R*_*minor*_ measurement is manually added as a column to the pandas dataframe storing PyMOL measurements with the column “R minor”.

#### Construction and calculation of *B*_*energetic*_

We defined a contact atom as a non-hydrogen atom of either the receptor or the peptide within 5Å of the binding partner in the peptide-receptor model since atoms within this distance cutoff are responsible for most protein-protein interactions^59^. We defined a clashing term as the number of non-hydrogen atoms in the peptide that are within 1Å of the receptor since this distance is smaller than the typical diameter of an atom and can cause a huge Van der Waals strain. To find the suitable weight for the clashing term, we estimated the relative energy scales using Lennard-Jones’ potential. We concluded that an order of magnitude of 10^3^ should be proper (Eq. 1). Since most interfaces between the engineered peptide and the receptor have up to a few hundred non-hydrogen atoms (tens of residues) in the interface, this heavy weight for the clashing atom practically sets the 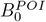 of any peptide with steric clashing against the receptor to 0. Thus, Eq. 1 is practically equivalent to:

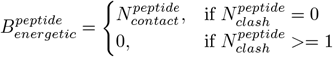

#### Construction and calculation of geometrical scores

The binding angle *θ* is defined as the angle between the vector from receptor center of mass to receptor anchor 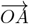 and the vector from receptor center of mass to peptide center of mass 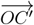 (Figure 2c. Note that the peptide center of mass *C*′ is usually very close to the deepest point *C*, and therefore point *C* and point *C*′ are undifferentiated in this schematic). A steep function is used to penalize inaccessible binding angles that are close to the anchor point:

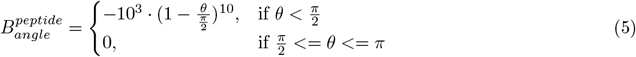

The definition of binding depth *d* is a simplification of previously defined travel depth ^60^: we first calculate the hydrodynamic radius of the hull of the receptor at the minor axis (*R*_*minor*_) using HullRad ^58^, and then take the difference of the distance between the “closest point on the peptide” to the receptor center and *R*_*minor*_. The ratio between the difference and *R*_*minor*_ is defined as the depth. In other words, binding depth 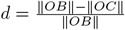 where ∥*OB*∥ is the minor axis radius (in Å) of the receptor hull when considering it as an ellipsoid (Figure 2c), and ∥*OC*∥ is the distance (in Å) between the center of mass of the receptor and the closest point on the peptide (Figure 2c). An odd polynomial function is used to construct the score to reflect both the positive effect of a deep binding pocket and the negative effect of a convex binding site:

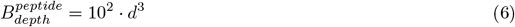

#### Calculation of scores for each peptide in a model

The total binding scores for each peptide in a model are calculated using Eqs. 1-4 in the main text.

#### Generation of the score matrix and a ranking

The total binding scores of a POI vs. a competitor across 10 replicate models are averaged to get 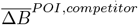 (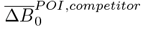 for APPRAISE 1.0, 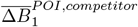 for APPRAISE 1.1, or 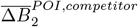 for APPRAISE 1.2). These averaged competition scores are then used to create a matrix and are plotted as a heatmap.

In the final score matrix, the POIs are ranked using a point-based round-robin tournament system^61^ to avoid the bias caused by individual competitions with unusually high scores. Briefly, each 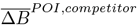 in the matrix is considered as the match result between a POI and a competitor. A POI gains 1 point for winning over each match and loses 1 point for losing each match. (In the cases when 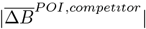 does not reach the threshold of *p <* 0.05 using a one-sample, two-sided, Student’s t test (degree of freedom=9), the match is called a tie, and the POI gets 0 points from the match.)

## Code Availability

All code is available in a GitHub repository:

github.com/GradinaruLab/APPRAISE

In addition, APPRAISE is made accessible through a web-based notebook interface using Google Colaboratory. The notebook can be found in the GitHub repository above or be directly accessed through the link:

tiny.cc/APPRAISE

The Colab-APPRAISE notebook includes pre-filled templates that can be used to demonstrate the workflow of APPRAISE. More demos can be found under the demo folder in the GitHub repository.

## Experimental Validations

### *In vitro* infectivity assay

HEK293T (ATCC, CRL-3216) cells were seeded in 6-well plates at 80% confluency and maintained in Dulbecco’s Modified Eagle Medium (DMEM) supplemented with 5% fetal bovine serum (FBS), 1% non-essential amino acids (NEAA), and 100 U/mL penicillin-streptomycin at 37°C in 5% CO2. Cells were transiently transfected with 2.53 μg plasmid DNA encoding an expression cassette for the LY6A receptor. The following day, receptor-expressing cells were transferred to black, clear-bottom 96-well plates at 20% confluency and maintained in FluoroBrite™ DMEM supplemented with 0.5% FBS, 1% NEAA, 100 U/mL penicillin-streptomycin, 1x GlutaMAX, and 15 μM HEPES at 37°C in 5% CO2. Engineered AAV variants packaging a CAG-mNeonGreen transgene were dosed in triplicate at 5E8 vg per well once the cells were attached. Plates were imaged 24 hours after AAV was introduced to cells with a Keyence BZ-X700 using a 4x objective and NucBlue™ Live ReadyProbes™ Reagent (Hoechst 33342) to autofocus each well.

### *In vivo* mouse experiment

For all the experiments performed in this study, the animals were randomly assigned, and the experimenters were not blinded while performing the experiments unless mentioned otherwise. All animal procedures in mice were approved by the California Institute of Technology Institutional Animal Care and Use Committee (IACUC), Caltech Office of Laboratory Animal Resources (OLAR), and were carried out in accordance with guidelines and regulations.

For the profiling of the novel AAVs in C57BL/6J mice (The Jackson Laboratory, 000664) and BALB/cJ mice (The Jackson Laboratory, 000651), the AAV vectors were injected intravenously via the retro-orbital route to 6-8 week-old adult mice at a dose of 3 × 10^11^ vg per mouse. Retro-orbital injections were performed as described previously ^62^. To harvest the tissues of interest after 3 weeks of expression, the mice were anesthetized with Euthasol (pentobarbital sodium and phenytoin sodium solution, Virbac AH) and transcardially perfused using 50 mL of 0.1 M phosphate-buffered saline (PBS) (pH 7.4), followed by 50 mL of 4% paraformaldehyde (PFA) in 0.1 M PBS. The organs were collected and post-fixed 24 h in 4% PFA at 4°C. Following this, the tissues were washed with 0.1 M PBS and stored in fresh PBS-azide (0.1 M PBS containing 0.05% sodium azide) at 4°C. Before imaging, the 100 μm tissue slices were cut using a Leica VT1000S. Brain images were acquired with a Zeiss LSM 880 confocal microscope using a Plan-Apochromat 10× 0.45 M27 (working distance 2.0 mm) objective. Zen Black 2.3 SP1 was used to process the images.

### Ethical approval

All animal procedures in mice were approved by the California Institute of Technology Institutional Animal Care and Use Committee (IACUC), Caltech Office of Laboratory Animal Resources (OLAR) and were carried out in accordance with guidelines and regulations.

## Materials Availability

The plasmid expressing the AAV-PHP.D capsid reported in this manuscript is deposited in Addgene (ID: 197055).

## Supplementary Information

### Supplementary Figures

**Fig. S1.**
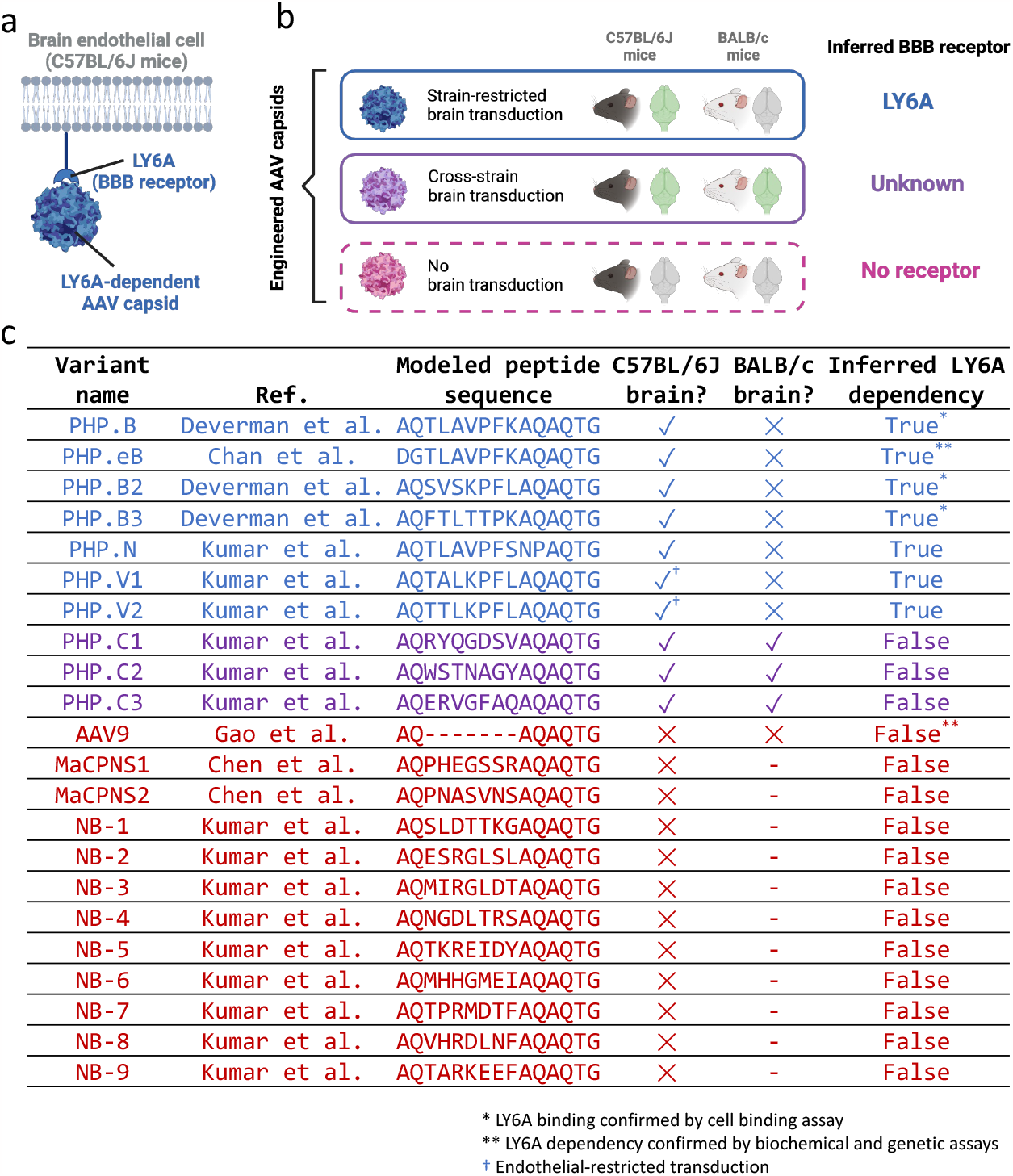
Prior experimental studies revealing the receptor dependency of some brain-transducing AAV variants. **a**, A schematic showing an AAV capsid binding to a blood-brain barrier (BBB) receptor that is only expressed at a high level on the endothelial cells of certain mouse strains ^37–39^. **b**, A schematic showing how we can infer whether an AAV capsid can use LY6A, a mouse BBB receptor, by characterizing its brain transduction across different strains. A capsid with strain-restricted brain transduction in C57BL/6J mice is likely LY6A-dependent, while a capsid with cross-strain brain transduction or which does not transduce the brain is not LY6A-dependent. **c**, A table summarizing all 22 capsids used in Figure 2 with their source literature, sequence, brain transduction profile, and inferred LY6A dependency.

**Fig. S2.**
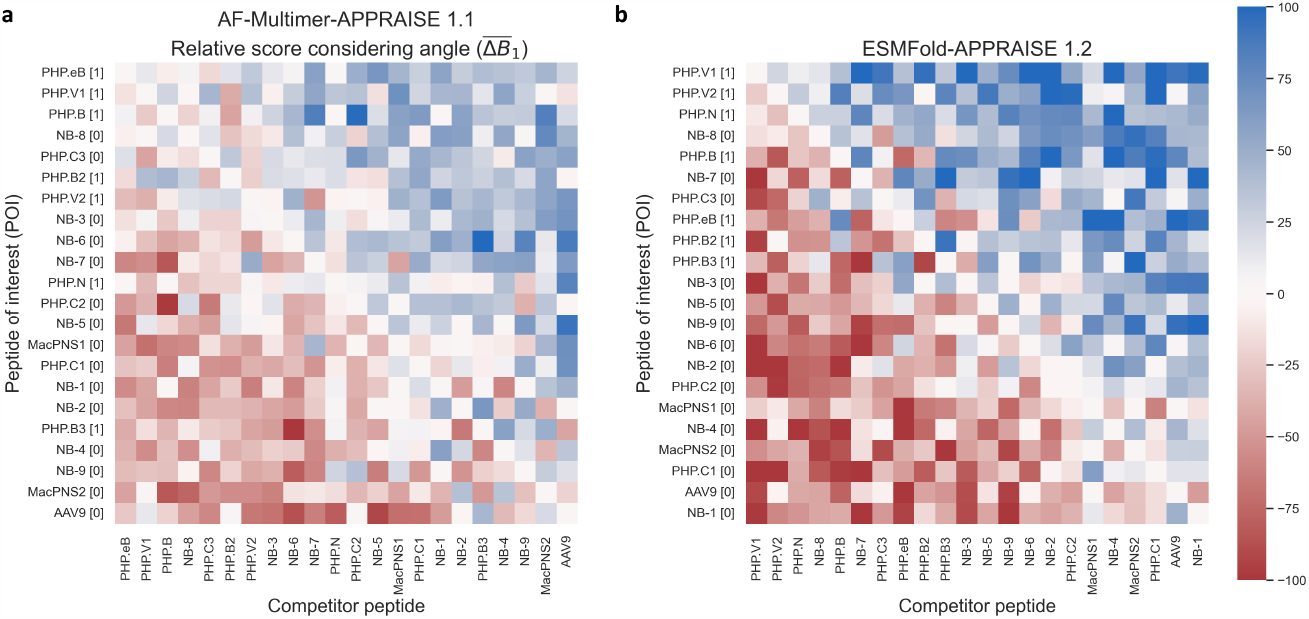
Heatmaps representing score matrices of AF-Multimer-APPRAISE 1.1 and ESMFold-APPRAISE 1.2.

**Fig. S3.**
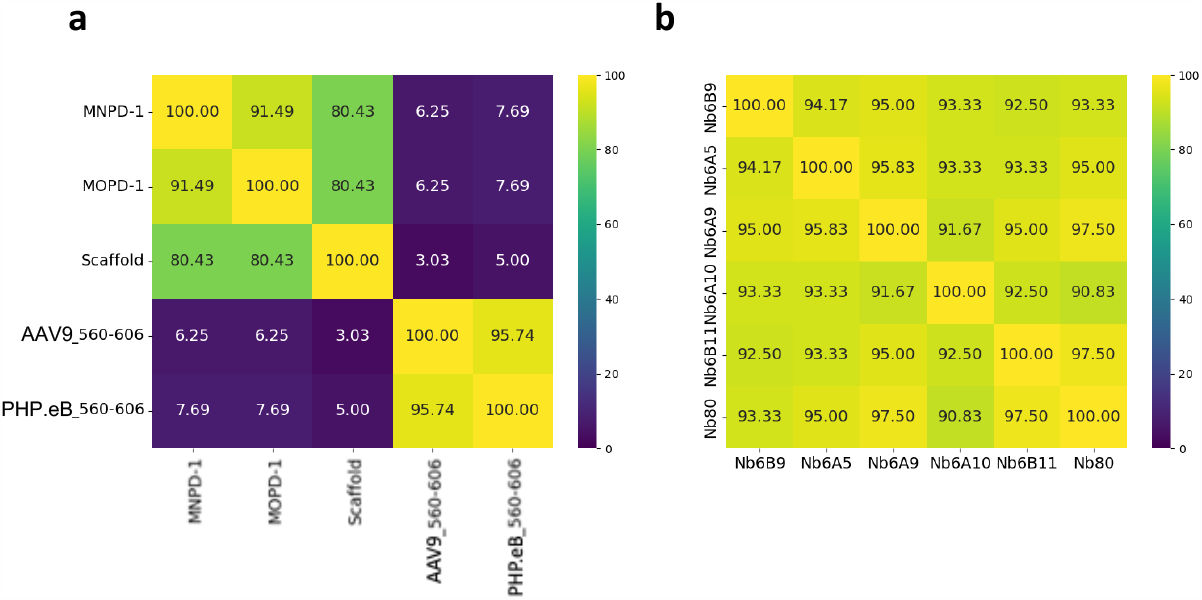
Sequence identity between some engineered proteins. Sequence identity matrix generated with Clustal Omega ^46^ 9 for **a)** PD-L1-binding peptides and **b)** *β*_2_ adrenergic receptor-binding nanobodies, demonstrating that these proteins share similar 0 sequences.

**Fig. S4.**
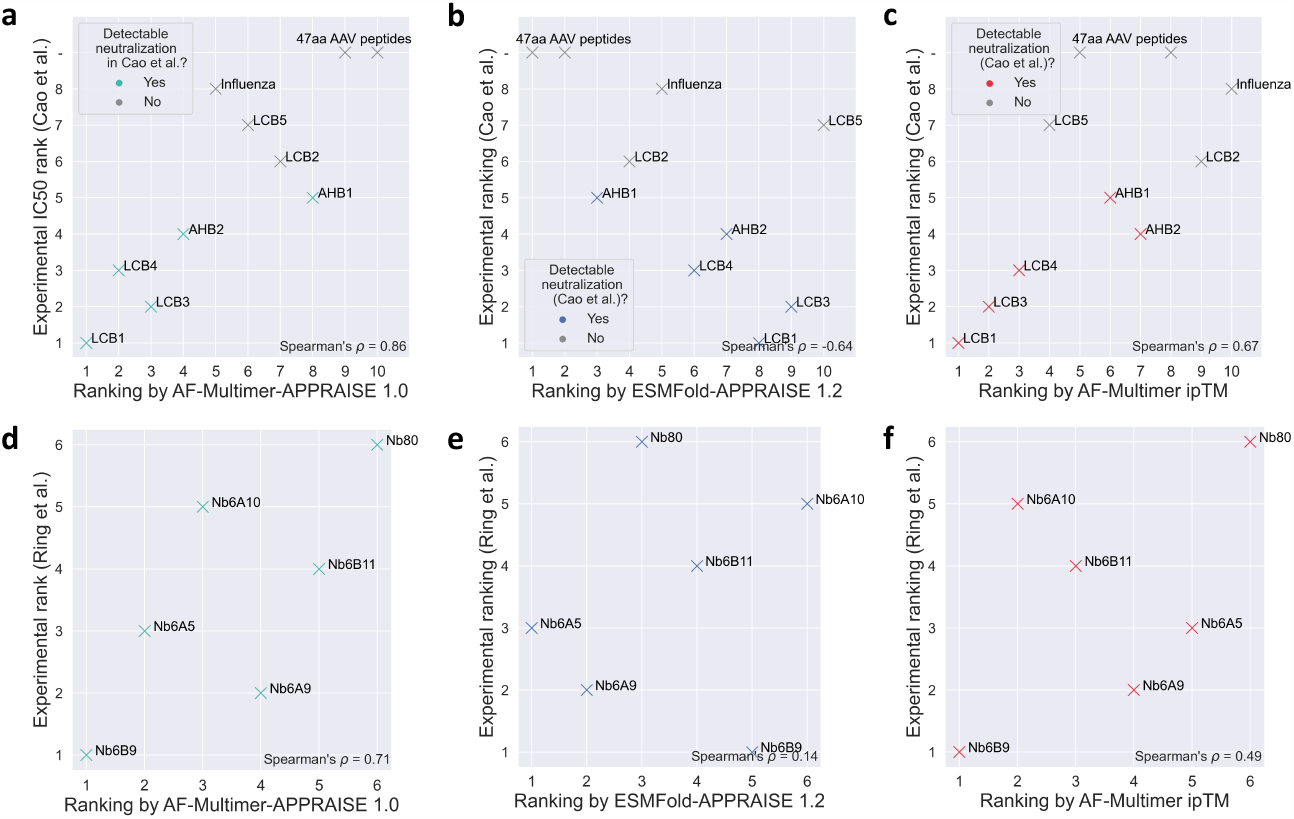
Ranking protein binders using alternative methods. Rankings of two groups of peptides analyzed in Figure 3g, j based on **a, d)** AF-Multimer-APPRAISE 1.0, **b, e)** ESMFold-APPRAISE 1.2, and **c, f)** interface pTM given by AlphaFold-Multimer.

**Fig. S5.**
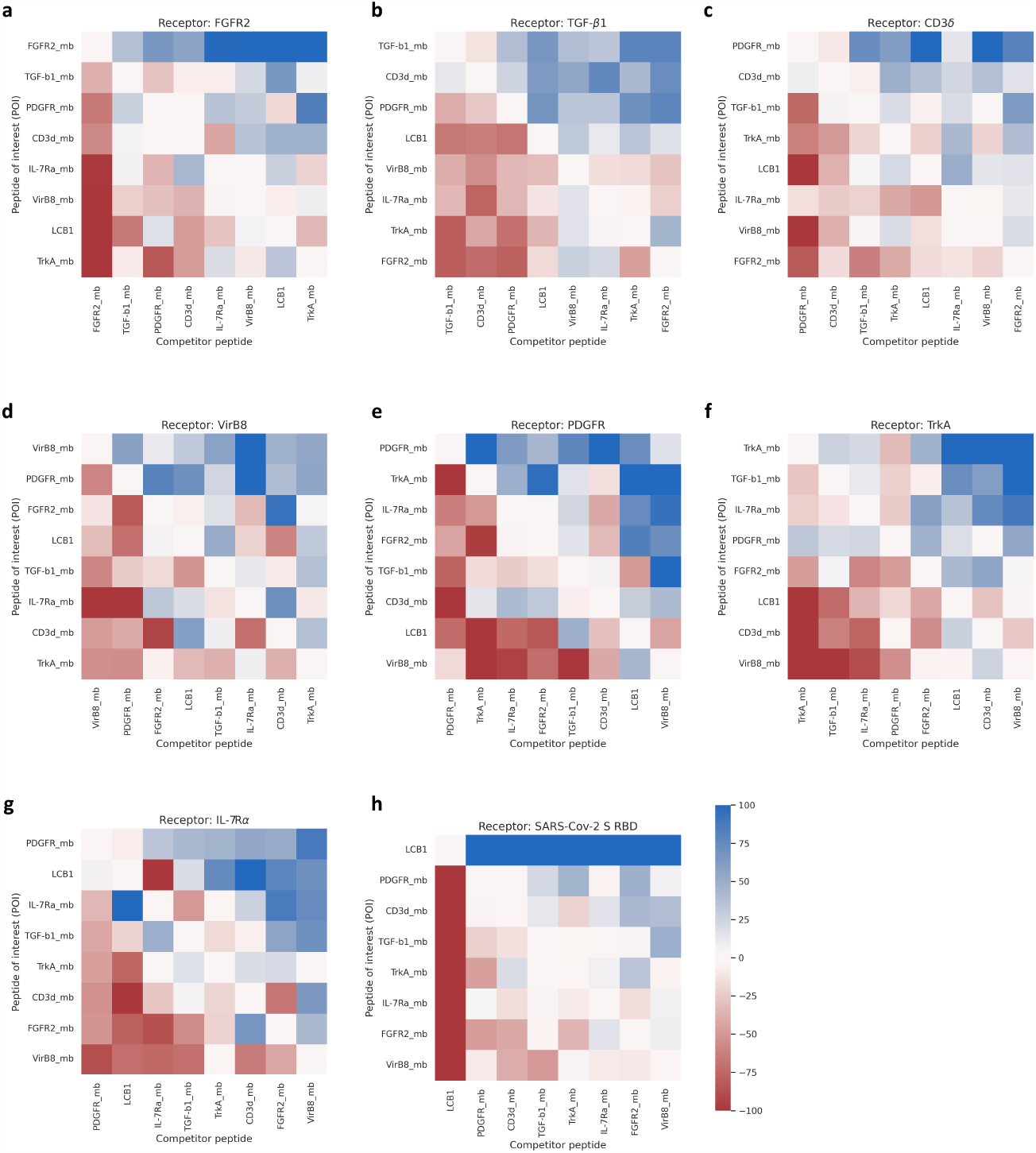
Score matrices for APPRAISE rankings of miniprotein binders with individual receptors. Detailed rankings of miniproteins with individual receptors summarized in Figure 3k using AF-Multimer-APPRAISE 1.2. The miniprotein sequences were from ^45^.

**Fig. S6.**
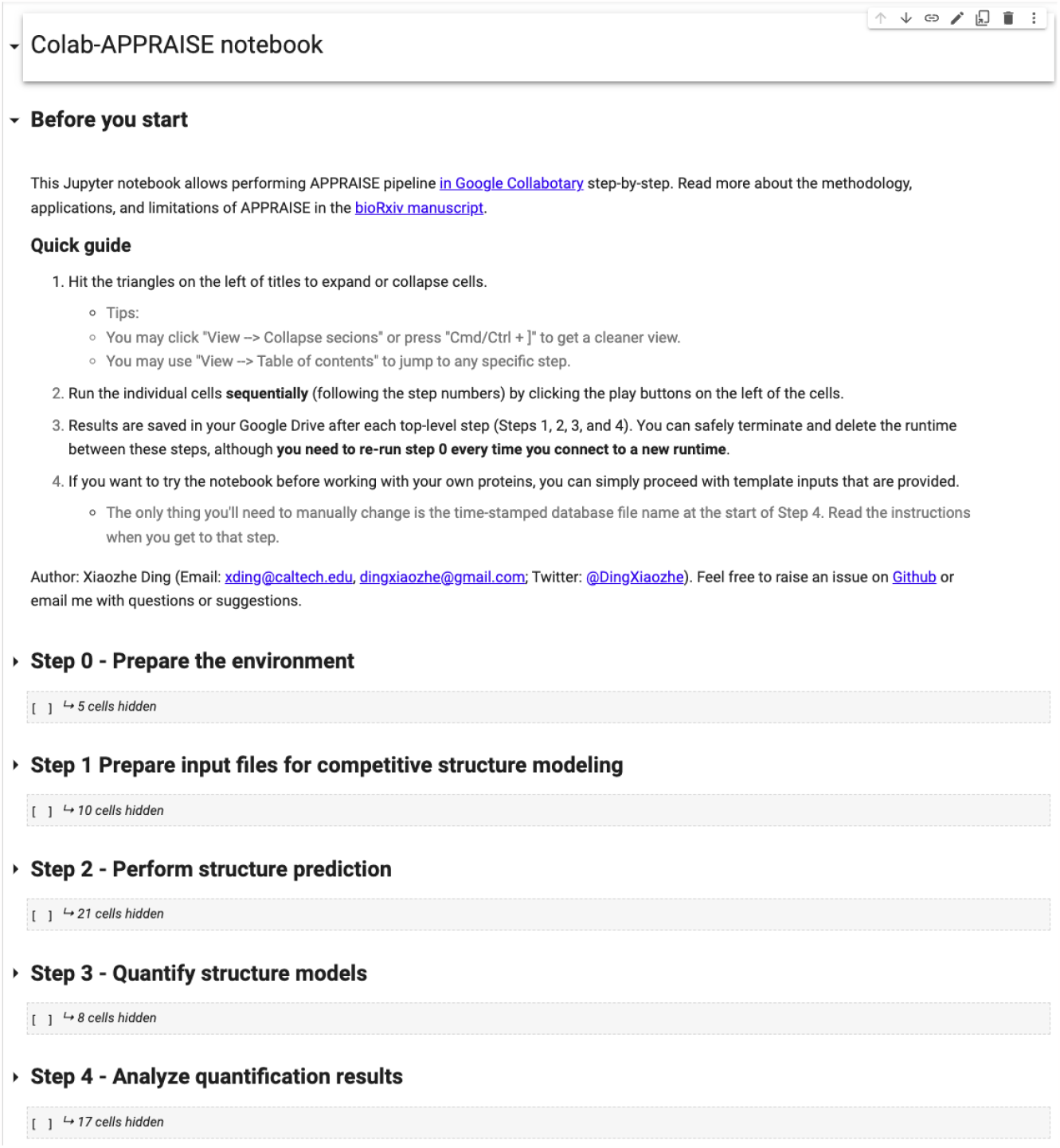
A screenshot of the interface of Colab-APPRAISE. APPRAISE can be easily accessed by running a web-based notebook on Google Colaboratory (https://tiny.cc/APPRAISE).

**Table S1.** Receptor sequences and parameters used for APPRAISE analysis (part 1 of 2)

| Receptor name | Organism | Uniprot accession ID | Domain used for modeling | Residue indices (start-end) | $D_{max}$ (Å) | Axial ratio | $R_{minor}$ (Å) | Anchor site | Sequence used for modeling |
| --- | --- | --- | --- | --- | --- | --- | --- | --- | --- |
| LY6A | Mus musculus (Mouse) | P05533 | Mature protein | 27-110 | 46.68 | 1.74 | 13.4 | C-term | LECYQCYGVPPFTSCPSITCYPDGVGVCTQEAIVDSQ<br>TRKVKWNLCLPICPPNIESMEILGTVKVVKTSCCQEDLC<br>NVAVP |
| PD-L1 | Homo sapiens (Human) | Q9NZQ7 | V domain | 18-132 | 46.50 | 1.51 | 15.4 | C-term | AFVTYVPKDLVYVEYGSNMTIECKFPVEKQLDLAALIVY<br>WEWEDKNIIQFVHGEECLKVQHSYRQARLLKQDLSLG<br>NAALQITDVKLQDAGVYRCMISYGGADYKRITVKVNA |
| Beta-2 adrenergic receptor | Homo sapiens (Human) | P07550 | TM1-TM7 | 29-342 | 93.8 | 2.23 | 21.0 | N-term | DEVVWVGIVMSLIVLAIIVFGNVLVITAIKFERLQTV<br>TNYFITSACADLVMLGAVVPPGAAHILMKMTFGNFWC<br>EFWTSIDVLCVTASITELCVIAVDRYFAITSPFKYQSL<br>TKNKARVILMWIVSGLSFLLPIQMHVYRATHQEAINE<br>YANETCCDFFTNQAYAIASSIVSFYVPLVIMVVFYSRVF<br>QEAQRQLQKIDKSEGRFHQNLQSQVEQDQRTGHGLRRSS<br>KFCLKEHKALKTGLIINGTFTLCWLPFFIVNIHVVIQDN<br>LIRKEVYILLNWIGYVNSGFNPLIYCRSPDFRIAFQELL<br>CL |
| Transferrin receptor 1 | Homo sapiens (Human) | P02786 | Ectodomain | 122-760 | 86.50 | 1.47 | 29.4 | N-term | LYWDDLKRLKSEKLDSTDFGTIKLLNENSVPREAGSQ<br>KDENLALYVENQFREFKLKSVWRQHFVKIQVKDSAQNS<br>VIIVDKNGRLVLYVENPGYVAYSKAATVTGKLVHANFG<br>TKKDFEDLYTPVNGSIVIVRAGKITFAEKVANAESLNAI<br>GVLIMYDQTKFPIVNAELSFHGAHLGTGDPYTPGPPSF<br>NHTQFPSSRSGLPNIPVQITISRAAAEKLFGNMEGDCPS<br>DWKTDSTCRMTSESKNVKLVTSNVLKEIKILNIFGVIK<br>GFVEPDHYVVVGAQRDAGPAAKSGVGTALLLKLQAMF<br>SDMVKDGFQPSRSIIFASWSAGDFGSGVATEWLEGYLS<br>SLHLKAFYIINLDKAVLGTNSFKVSPPLYTLIEKTMQ<br>NVKHPVTGQFLYQDSNWSKVEKLTLDNAFPFLAYSIGI<br>PAVSFCFCEDTDYPLYGTTMDTYKELIERIPELNKVARA<br>AAEVAGQFVILKTHDVELNLDYERYNSQLLSFVRDLNQY<br>RADIKEMGLSLQWLYSARGDFFRATSLRTDFFGNAEKTD<br>RFVMMKLNDRVMRVEYHFLSPYVPSKESPRHVFWSGS<br>HTLPALLENLKLKQNGNAFNETFRNGLALATWTIQGA<br>ANALSGDVWDIDNEF |
| Spike | SARS-CoV-2 | PODTC2 | RBD | 331-529 | 67.93 | 1.87 | 18.2 | C-term | NITNLCPGEVFNATRFASVYAWNKRISNCVADSVLY<br>NSASFSTFKCYGVSPTKLMDLCTNIVYADSFVRIGDEV<br>QIAPQGTGKIADYNYKLPDDFTGCVIAWNSNLDKSVGG<br>NYYNLYRLFRKSNLKPFDISTEIVQAGSTPCNGVEGF<br>NCYFPLQSYGFQPTNGVGYQPYRVVVLSFELLHAPATVC<br>GPKK |

**Table S2.** Receptor sequences and parameters used for APPRAISE analysis (part 2 of 2)

| Receptor name | PDB ID | Chain ID | Regions used for modeling | Construct length | $D_{max}$ (Å) | Axial ratio | $R_{minor}$ (Å) | Anchor site | Sequence used for modeling |
| --- | --- | --- | --- | --- | --- | --- | --- | --- | --- |
| FGFR2 | 1DJS | A | Structured region | 216 | 105.22 | 3.06 | 17.19 | C-term | TLEPEGAPYWTNTEKMEKRLHAVPAANTVKFRCPAGGNP<br>MPTMRWLKNGKEFKQHRIGGYKVRNQHSIMESVVPVS<br>DKGNYTCVVENEYGSINHLYDLVDVERSPHRILQAGLP<br>ANASTVGGDVEFVCKVYSDAQPHIQIKHVEKNGSKYG<br>PDGLPYLVKLAAGVNTDKIEIYLYIRNVTFEDAGEYT<br>CLAGNSIGISFSAWLTPLA |
| TGF-β1 | 3KFD | A | Structured region | 112 | 65.88 | 2.4 | 13.73 | C-term | ALDTNYCFSSTEKNCCVRQLYIDFRKDLGWKWIHEPKGY<br>HANFCLGCPYIWSLDTQYSKVLALYQHNHPGASAAPCC<br>VPQALEPLPIVYVGRKPKVEQLSNMIVRSCKCS |
| CD3δ | 1XIW | B | Structured region | 74 | 48.7 | 2.06 | 11.82 | C-term | MKIPIEELEDREVFNCTSIITWEGTVGTLSDITRLDL<br>GKRILDPRIYRCNGTDIYKDKESTVQVHYRMCQS |
| VirB8 | 403V | A | Cytoplasmic domain | 143 | 56.27 | 1.62 | 17.37 | C-term | ANPYISVANIMLQNVKQREKYNYDTLKEQFTFIKNAST<br>SIVYMQFANFMINDSLSPVIRYQKLYRSINIIISINNI<br>NNNEATVTFESLAQNNTGEILENMLEAKIGFIMDSIST<br>STLHNMPEHFIVTSYKLLRLNKNQ |
| PDGFR | 3MJG | C | Ig-like C2-type domains 2&3 | 202 | 86.04 | 2.33 | 18.46 | C-term | DERKRLYIFVDPPTVGFPLNDAEELFIFLTEITEITIPC<br>RVTDPQLVVTLEHKKGDVALPVYDHRQGFSGIFEDRSY<br>ICKTTIGDREVDSDAYVYVRLQVSSINVSNAVQTVVRQ<br>GENITLMCIVIGNEVVMFETYPKESGRLEVPTDFLL<br>DMPYHISILHPSAELEDSCYTTCNVTESVNDHQDEKA<br>INITVVE |
| TrkA | 2IFG | A | Ig-like C2-type domain 2 | 100 | 50.52 | 1.8 | 14.03 | C-term | SFPASVQLHTAVEMHHWCIPFSVDGQAPSLRLWLFNGSV<br>LNETSIFTFLEPAANETVRHGLRLMQPTHVNNNGHYT<br>LLAANPFGQASASIMAFMDNP |
| IL-7Rα | 3DI3 | B | Structured region | 193 | 70.65 | 2.07 | 17.07 | C-term | DYSCYCYQLVNGSQHSLTCAFEDPDVNTWLEFEICG<br>ALVEVKCLNFRKLQEIYFIETKKLLIGKSNICVKVGEK<br>SLTCKKIDLTITVKEPAFDLSVYVREGANDFVVTFTS<br>HLQKVKVLMHDVAYRAQEKDENKWNHNLSSKTLTLQG<br>RKLQPAAMYIEKVRISIPDHYFKGFWSWSPSYFRTP |

**Table S3.**
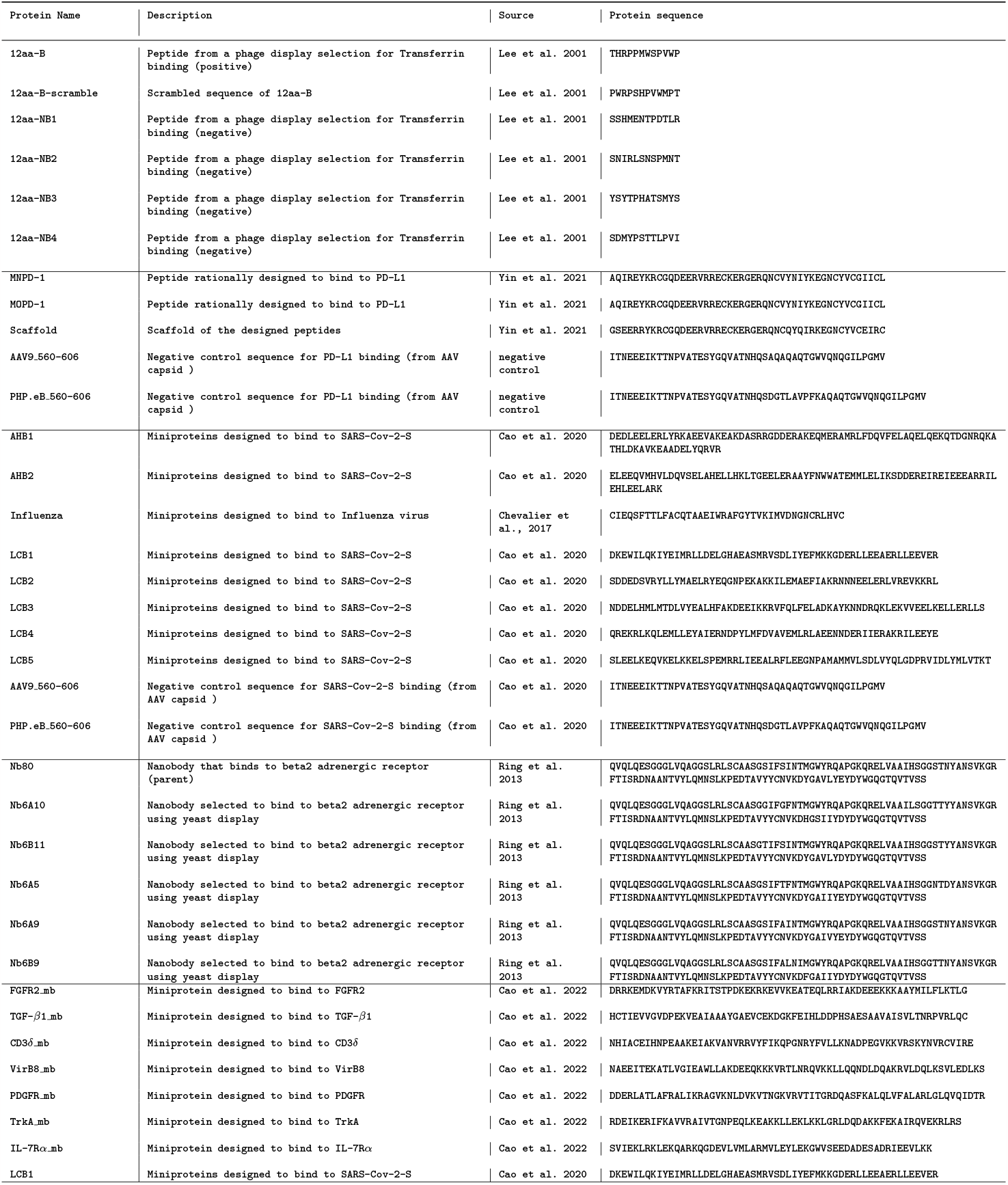
Sequences of engineered proteins used in APPRAISE tests in Figure 3.

**Table S4.** Peptides used for *in silico* screening.

| Peptide name | Peptide sequence (residues 587-594 in VP1) | Original source |
| --- | --- | --- |
| AAV9 | AQ-----AQAQTG | Gao et al., 2002 |
| PHP-B | AQTLAVPFKAQAQTG | Deverman et al. 2016 |
| PHP-D | AQWKMMGLQAQAQTG | Unpublished *in vivo* selection |
| SRK-1 | AQLYHGGSTAQAQTG | Kumar et al. 2020 |
| SRK-2 | AQNNSVRQLAQAQTG | Kumar et al. 2020 |
| SRK-3 | AQVNSTRNVAQAQTG | Kumar et al. 2020 |
| SRK-4 | AQGNMTKFTAQAQTG | Kumar et al. 2020 |
| SRK-5 | AQTAIQPPKAQAQTG | Kumar et al. 2020 |
| SRK-6 | AQITTDQPPFAQAQTG | Kumar et al. 2020 |
| SRK-7 | AQDTANTARAQAQTG | Kumar et al. 2020 |
| SRK-8 | AQTHDAQAWAQAQTG | Kumar et al. 2020 |
| SRK-9 | AQQPLAEAAQAQTG | Kumar et al. 2020 |
| SRK-10 | AQTALANQKAQAQTG | Kumar et al. 2020 |
| SRK-11 | AQTGTERLSAQAQTG | Kumar et al. 2020 |
| SRK-12 | AQNGVVTQSKAQAQTG | Kumar et al. 2020 |
| SRK-13 | AQWTEQRLVAQAQTG | Kumar et al. 2020 |
| SRK-14 | AQDTGLNNRAQAQTG | Kumar et al. 2020 |
| SRK-15 | AQPLPPTSIAQAQTG | Kumar et al. 2020 |
| SRK-16 | AQSDPGKFMAQAQTG | Kumar et al. 2020 |
| SRK-17 | AQITTMGTMLAQAQTG | Kumar et al. 2020 |
| SRK-18 | AQKQTQDSSAQAQTG | Kumar et al. 2020 |
| SRK-19 | AQLAHNSALQAQAQTG | Kumar et al. 2020 |
| SRK-20 | AQVVPSTYRAQAQTG | Kumar et al. 2020 |
| SRK-21 | AQFRHLTGAQAQTG | Kumar et al. 2020 |
| SRK-22 | AQSANLLSSAQAQTG | Kumar et al. 2020 |
| SRK-23 | AQFSNTHALAQAQTG | Kumar et al. 2020 |
| SRK-24 | AQFNSKLQLAQAQTG | Kumar et al. 2020 |
| SRK-25 | AQFNTNISAAQAQTG | Kumar et al. 2020 |
| SRK-26 | AQYVPVLKQAQAQTG | Kumar et al. 2020 |
| SRK-27 | AQHVNHMMAQAQAQTG | Kumar et al. 2020 |
| SRK-28 | AQIVSNQMSAQAQTG | Kumar et al. 2020 |
| SRK-29 | AQPRPERMYAQAQTG | Kumar et al. 2020 |
| SRK-30 | AQNMKIQHVAQAQTG | Kumar et al. 2020 |
| SRK-31 | AQNTNVPPAMAQAQTG | Kumar et al. 2020 |
| SRK-32 | AQSAQLRSSAQAQTG | Kumar et al. 2020 |
| SRK-33 | AQSHHEQVSAQAQTG | Kumar et al. 2020 |
| SRK-34 | AQGAATGHLTAQAQTG | Kumar et al. 2020 |
| SRK-35 | AQHNLRDSIAQAQTG | Kumar et al. 2020 |
| SRK-36 | AQPGTGSFKAQAQTG | Kumar et al. 2020 |
| SRK-37 | AQSPPPVQGLAQAQTG | Kumar et al. 2020 |
| SRK-38 | AQTLVNAIHAQAQTG | Kumar et al. 2020 |
| SRK-39 | AQLGDDITGFAQAQTG | Kumar et al. 2020 |
| SRK-40 | AQGFNSMKFAQAQTG | Kumar et al. 2020 |
| SRK-41 | AQSNGLNGLAQAQTG | Kumar et al. 2020 |
| SRK-42 | AQVRIPGALAQAQTG | Kumar et al. 2020 |
| SRK-43 | AQDMGTDNLQAQAQTG | Kumar et al. 2020 |
| SRK-44 | AQNYATKSQAQAQTG | Kumar et al. 2020 |
| SRK-45 | AQSVTTSHVAQAQTG | Kumar et al. 2020 |
| SRK-46 | AQTSGTGDLAQAQTG | Kumar et al. 2020 |
| SRK-47 | AQARTAHGYAQAQTG | Kumar et al. 2020 |
| SRK-48 | AQHSANMSKAQAQTG | Kumar et al. 2020 |
| SRK-49 | AQHDERANMAQAQTG | Kumar et al. 2020 |
| SRK-50 | AQNNFNASLAQAQTG | Kumar et al. 2020 |
| SRK-51 | AQSAASLVSHAQAQTG | Kumar et al. 2020 |
| SRK-52 | AQAPRIDNAAQAQTG | Kumar et al. 2020 |
| SRK-53 | AQLTSSNALAQAQTG | Kumar et al. 2020 |
| SRK-54 | AQTLNSTRAAQAQTG | Kumar et al. 2020 |
| SRK-55 | AQSGTCRQQAQAQTG | Kumar et al. 2020 |
| SRK-56 | AQKTTLASGAQAQTG | Kumar et al. 2020 |
| SRK-57 | AQMRVNTTEAQAQTG | Kumar et al. 2020 |
| SRK-58 | AQFETLHKTAQAQTG | Kumar et al. 2020 |
| SRK-59 | AQTQHRFEMAQAQTG | Kumar et al. 2020 |
| SRK-60 | AQHTAEKAPAQAQTG | Kumar et al. 2020 |
| SRK-61 | AQNHMVRELAQAQTG | Kumar et al. 2020 |
| SRK-62 | AQRFPQSSAQAQTG | Kumar et al. 2020 |
| SRK-63 | AQRSVANVPAQAQTG | Kumar et al. 2020 |
| SRK-64 | AQVFQATRTAQAQTG | Kumar et al. 2020 |
| SRK-65 | AQEQRTPSPAQAQTG | Kumar et al. 2020 |
| SRK-66 | AQGSSTASLAQAQTG | Kumar et al. 2020 |
| SRK-67 | AQQVPHLHSAQAQTG | Kumar et al. 2020 |
| SRK-68 | AQPSQPYYKAQAQTG | Kumar et al. 2020 |
| SRK-69 | AQTHTRDQGAQAQTG | Kumar et al. 2020 |
| SRK-70 | AQINPGITLAQAQTG | Kumar et al. 2020 |
| SRK-71 | AQLQPTKSSAQAQTG | Kumar et al. 2020 |
| SRK-72 | AQQDAKVTTAQAQTG | Kumar et al. 2020 |
| SRK-73 | AQGASTHNAQAQTG | Kumar et al. 2020 |
| SRK-74 | AQIPVSIQAQAQTG | Kumar et al. 2020 |
| SRK-75 | AQVTSAHPVAQAQTG | Kumar et al. 2020 |
| SRK-76 | AQTASLLASAQAQTG | Kumar et al. 2020 |
| SRK-77 | AQDRGTRTVQAQTG | Kumar et al. 2020 |
| SRK-78 | AQTAYLEVKAQAQTG | Kumar et al. 2020 |
| SRK-79 | AQATTQMSSAQAQTG | Kumar et al. 2020 |
| SRK-80 | AQKYDASQSAQAQTG | Kumar et al. 2020 |
| SRK-81 | AQTGTSHLHAQAQTG | Kumar et al. 2020 |
| SRK-82 | AQTMTPSGIAQAQTG | Kumar et al. 2020 |
| SRK-83 | AQTPSSSGNAQAQTG | Kumar et al. 2020 |
| SRK-84 | AQKDVVNSNAQAQTG | Kumar et al. 2020 |
| SRK-85 | AQRSPATMLAQAQTG | Kumar et al. 2020 |
| SRK-86 | AQYDQKSLAQAQTG | Kumar et al. 2020 |
| SRK-87 | AQMGARNLPAQAQTG | Kumar et al. 2020 |
| SRK-88 | AQLPISATEAQAQTG | Kumar et al. 2020 |
| SRK-89 | AQTRHTSLTAQAQTG | Kumar et al. 2020 |
| SRK-90 | AQNKLTANGAQAQTG | Kumar et al. 2020 |
| SRK-91 | AQNGDSHSHAQAQTG | Kumar et al. 2020 |
| SRK-92 | AQVRIDMDMAQAQTG | Kumar et al. 2020 |
| SRK-93 | AQSVSTPRGAQAQTG | Kumar et al. 2020 |
| SRK-94 | AQVSRQFEPQAQAQTG | Kumar et al. 2020 |
| SRK-95 | AQSANNVRGAQAQTG | Kumar et al. 2020 |
| SRK-96 | AQIGTKSTNAQAQTG | Kumar et al. 2020 |
| SRK-97 | AQGSSELRGAQAQTG | Kumar et al. 2020 |

## Acknowledgments

The authors thank Elisha Mackey, Zhe Qu, and Pat Anguiano for administrative assistance, Catherine Oikonomou for help with manuscript editing, and Sripriya R. Kumar, Seongmin Jang, Jimin Park, and Changfan Lin for helpful discussions. Schematics in this manuscript were created with BioRender.com. The study was funded by an NIH Director’s Pioneer Award DP1OD025535 (to V.G.).

## Declarations

### Declaration of Generative AI and AI-assisted technologies in the writing process

During the preparation of this work, the authors used Stork Writing Assistant in order to improve grammar and writing clarity. After using this tool/service, the authors reviewed and edited the content as needed and take full responsibility for the content of the publication.

### Ethics approval

All animal procedures in mice were approved by the California Institute of Technology Institutional Animal Care and Use Committee (IACUC), Caltech Office of Laboratory Animal Resources (OLAR), and were carried out in accordance with guidelines and regulations.

### Authors’ contributions

X.D. and V.G. conceived the project. X.D. developed the APPRAISE method and applied APPRAISE to predict the receptor binders. X.C. identified PHP.D through directed evolution and characterized PHP.D *in vivo*. E.E.S. and T.F.S. designed and performed *in vitro* infectivity assays.

